# Rapid promoter evolution of male accessory gland genes is accompanied by divergent expression in closely related *Drosophila* species

**DOI:** 10.1101/2024.10.14.618274

**Authors:** David W. J. McQuarrie, Frannie H. S. Stephens, Alexander D. Ferguson, Roland Arnold, Alberto Civetta, Matthias Soller

## Abstract

Seminal fluid proteins (Sfps) are essential for reproductive success and evolve fast, possibly driven by post-copulatory sexual selection (PCSS) originating from sperm competition and cryptic female choice. Counterintuitively, however, the coding region only in few Sfps evolves adaptively. Hence, additional genomic and functional factors must play a role in Sfp evolution independent of the protein coding region. To shed light on drivers of Sfp evolution we focus on those Sfps predominantly expressed in male accessory glands, because this allows examination of their evolution in the tissue which produces the majority of Sfps. Moreover, accessory glands develop normally in hybrids in contrast to testis allowing to control for changes in cellular environment arising during speciation. Here, we discover that Acp promoters contain hot spots for rapid evolution from accumulation of sequence changes and insertions/deletions (indels). We further show that changes in promoter sequences are accompanied by gene expression divergence among closely related *Drosophila* species. We then validate these observations in *Drosophila* hybrids to show that species-specific expression divergence of Acps with rapidly evolving promoters are maintained in hybrids for some Acps, while others show dominance of one allele, a phenomenon termed transvection. These results indicate that *cis*-regulatory evolution, rather than genome background variation, drives Acp expression changes and promotes their rapid evolution.

## Background

During copulation, sperm together with seminal fluid is transferred from males to females. Seminal fluid proteins (Sfps) have a range of vital functions, including regulation of sperm storage, sperm survival and fertilization, but also include complex roles in sperm competition [1–8]. Moreover, Sfps induce physiological and behavioural changes in females after mating which are essential for reproductive success [5, 9]. Most Sfps are products of the male reproductive tissue the accessory glands, called accessory gland proteins (Acps) [10]. A key regulator of the female post-mating response in *Drosophila melanogaster* is sex-peptide (SP), that upon transfer during mating enters the haemolymph and passes the blood brain barrier to target command neurons in the brain that reduce remating and induce oviposition [11, 12]. In addition, SP regulates many other aspects of the female post-mating response including increased egg production, feeding, a change in food choice, sleep, memory, constipation, midgut morphology, and stimulation of the immune system [13–22].

In general, reproductive genes evolve more rapidly than non-reproductive genes. Sfps are some of the fastest-evolving genes with many lacking detectable similarity or orthologs between closely related species [1, 5, 23–29]. Fast evolution of Sfps has to a large degree been attributed to post-copulatory selection (PCSS) originating from sperm competition and cryptic female choice due to widespread polyandry [1, 4, 30–32]. Surprisingly, however, coding regions of only a few Sfp genes are under positive selection in *Drosophila* [28, 33–38]. Rapid coding sequence evolution between of most Sfps between *Drosophila melanogaster* and *D. simulans* has been shown to occur through relaxed, allowing increased genetic variation within and between species [24].

Of note, expression of Sfp genes rapidly diverges between related species [39–44]. These observations support that divergence in gene expression between species can lead to phenotypic changes among species without altering the amino acid sequence [39, 45, 46]. Consequently, these expression changes in Sfp genes could influence female post-mating fitness and be selectively favoured for driving adaptive processes. In fact, a recent survey of selection acting upon expression of Sfp genes uncovered enrichment for both stabilizing and directional selection for Sfps with reproductive-tissue-specific expression [44]. However, how changes in gene expression are implemented into the gene structure to drive emergence of species-specific adaptations is poorly understood.

Divergence due to species-specific adaptations can cause hybrid incompatibilities manifested in sterility or lethality in hybrids of closely related species [47]. Unexpectedly, nuclear pore complex (NPC) genes, required for selective transport of cargoes between the nucleus and cytoplasm, are prominently represented among genes involved in speciation [48–50]. Interestingly, a promoter deletion allele in NPC gene *Nup54* has been identified that results in SP insensitivity from defects in neuronal wiring [51]. Since the *Nup54* promoter accumulates indels and base changes rapidly, a critical role for gene regulatory elements at the onset of speciation is indicated as a result of sexual conflict [51]. Moreover, systematic analysis of NPC promoters further revealed that rapid evolution of NPC promoters is a general feature of this class of genes [52]. This feature is further shared between germline genes including piRNA pathway factors that are regulated by the NPC [53, 54]. Since piRNAs are key to supress mobilisation of transposons [55, 56], it is feasible to envision increased mutagenicity from transposon insertions into promoter regions, because they harbour open chromatin and are accessible to transposon insertions.

To specifically address whether rapid evolution of promoters is a key feature in the process of speciation, we focused on genes coding for Acps because they constitute the majority of Sfps, are expressed in a single tissue consisting of only two cell types and expression of these genes can be analysed in whole genome RNA-seq datasets of closely related species and hybrids [10, 57]. Our analysis reveals that promoters of Acps evolve fast, in contrast to their coding regions, which do not show rapid evolution among closely related species. Moreover, these changes in promoters are paralleled by changes in expression between closely related species. While some Acp expression changes are maintained in hybrids, others show dominance for one species. Such dominance of one promoter allele is termed transvection and is a result of chromosome pairing [58]. Hence, we can conclude that promoter changes and not genomic background is the main driver to promote changes in expression suggesting that promoter evolution is a major driver for the rise of new species.

## Results

### Promoters of accessory gland genes are hot spots for accumulation of indels and base changes

To analyse promoter evolution of Acps we first generated a list of 155 bona fide genes predominantly expressed in male accessory glands from annotated gene expression data [1, 23, 59–61] (Supplementary Dataset 1). We then used PhyloP27way data, a conservation score based on 27 insect species at each nucleotide position, to generate comparative metagene plots of regions 1000 nucleotides upstream and 300 nucleotides downstream of transcription start sites (TSS) of Acp genes and compared them to all other genes in the *D. melanogaster* genome (Fig. 1A). Quantification of the metagene plot values and analysis of mutation hotpots between the Acp genes and all other genes revealed that Acps accumulate mutations more rapidly in their promoters compared to the genome (Fig. 1A-C). In addition, accumulation of mutations was also observed in 5′ untranslated regions (UTRs) of Acps, in contrast to the 5′UTRs of NPC genes and piRNA factors [52, 53]. Because Acp genes are generally small and compact genes, regulatory elements could possibly be located in 5′UTRs (Supplementary Dataset 1, Fig. 1A-C).

**Fig. 1:**
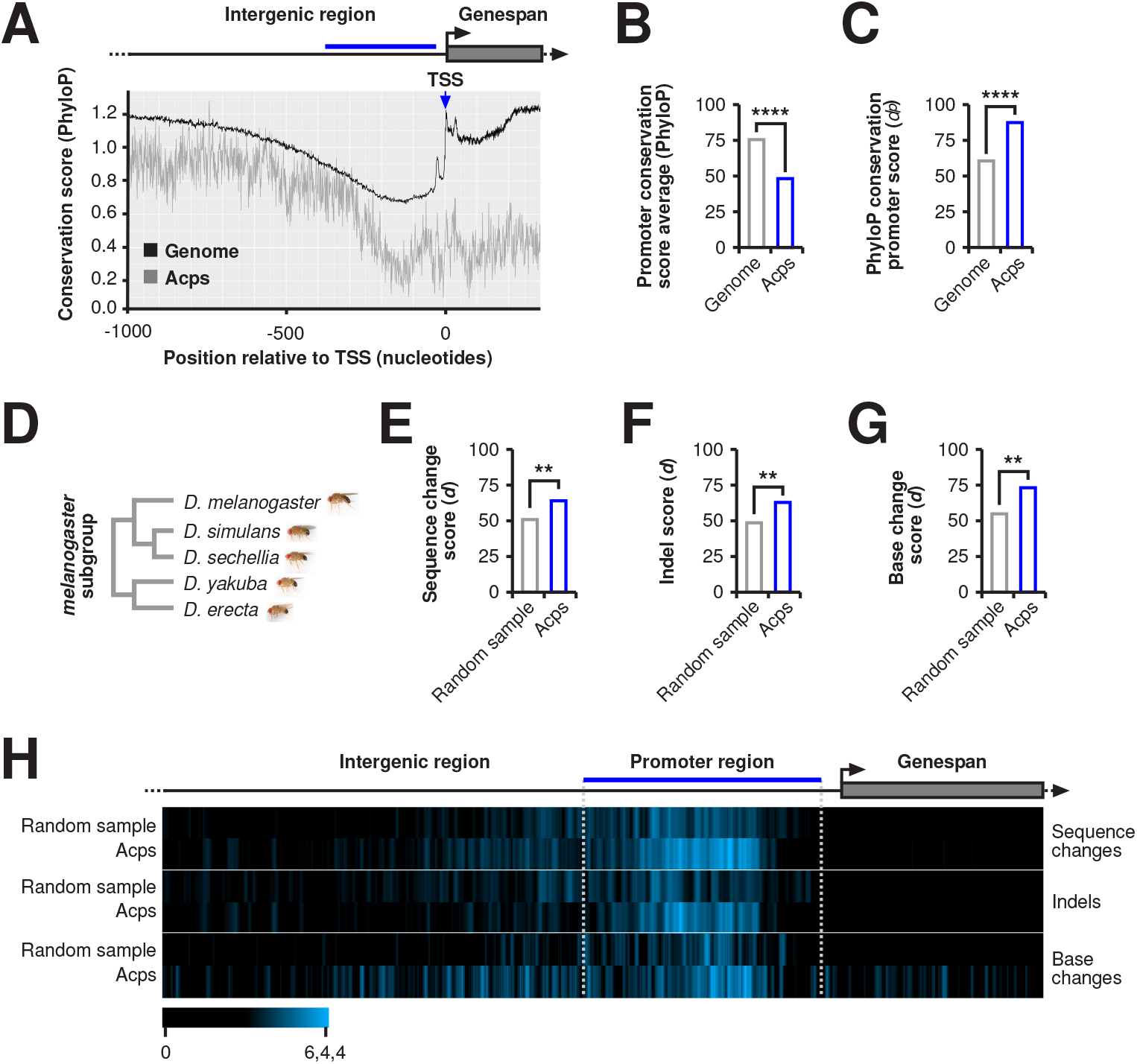
Promoters of accessory gland-specific Acp genes are hot spots for indel and base change accumulation. A) Metagene plot of average PhyloP27way conservations scores between Acps (grey) and all other genes in the *Drosophila* genome (black). The blue line indicates the analysed 350 nucleotide promoter region. B and C) PhyloP27way conservation score averages (B) and PhyloP27way conservation promoter *d^P^* scores (C) for the 350 nucleotide promoter regions of Acp genes compared to all genes in the *Drosophila* genome. Statistically significant differences from unpaired student t-tests (B) and non-parametric chi-squared tests (C) are indicated by asterisks (**** p≤0.0001). D) Phylogenetic tree of the *melanogaster* subgroup (*D. melanogaster*, *D. simulans*, *D. sechellia*, *D. yakuba*, and *D. erecta*) of *Drosophila* which were analysed in this study. E-G) Sequence change quantification (*d*) for Acps (blue) compared to a random sample of genes (grey) between the *melanogaster* subgroup members (D). Statistically significant differences from non-parametric chi-squared tests are indicated by asterisks (** p≤0.01). H) Heatmaps indicating a sliding window score of sequence change, indel, or base change accumulation -1000/+300 nucleotides from the TSS.

To understand whether promoter regions evolve rapidly between closely related *Drosophila* species, we focussed on the *melanogaster* subgroup which is made up of five species (*D. melanogaster, D. simulans, D. sechellia, D. yakuba,* and *D. erecta*) (Fig. 1D). We quantified accumulation of sequence changes in promoters for insertions and deletions (indels), and base changes in Acps and compared them to an equal number of random sampled genes (Fig. 1E-H). Our analysis confirmed that, compared to control genes, promoters of Acps evolve rapidly among closely related species by accumulating both indels and base changes.

### Rapidly diverging expression of Acp genes among closely related *Drosophila* species correlates with accumulation of mutations in promoters

To determine whether changes in promoter regions of Acp genes correlate with divergent gene expression, we examined differential gene expression between three pairs of closely related species in the *melanogaster* subgroup (*D. melanogaster* with *D. simulans*, *D. melanogaster* with *D. yakuba*, and *D. simulans* with *D. yakuba*). In this analysis we found that expression of Acp genes diverges more compared to all non-Acp genes and the random sample genes for the *D. melanogaster* versus *D. simulans* comparison (Fig. 2A). In addition, as a negative control we analysed highly conserved ribosomal component gene expression which showed significantly less divergence compared to the Acp genes for all comparisons (Fig. 2A). Consistent with a model for rapid evolution of changes in Acp gene expression, the distribution of expression changes is skewed for Acps but not for the random control sample (Fig. 2B).

**Fig. 2:**
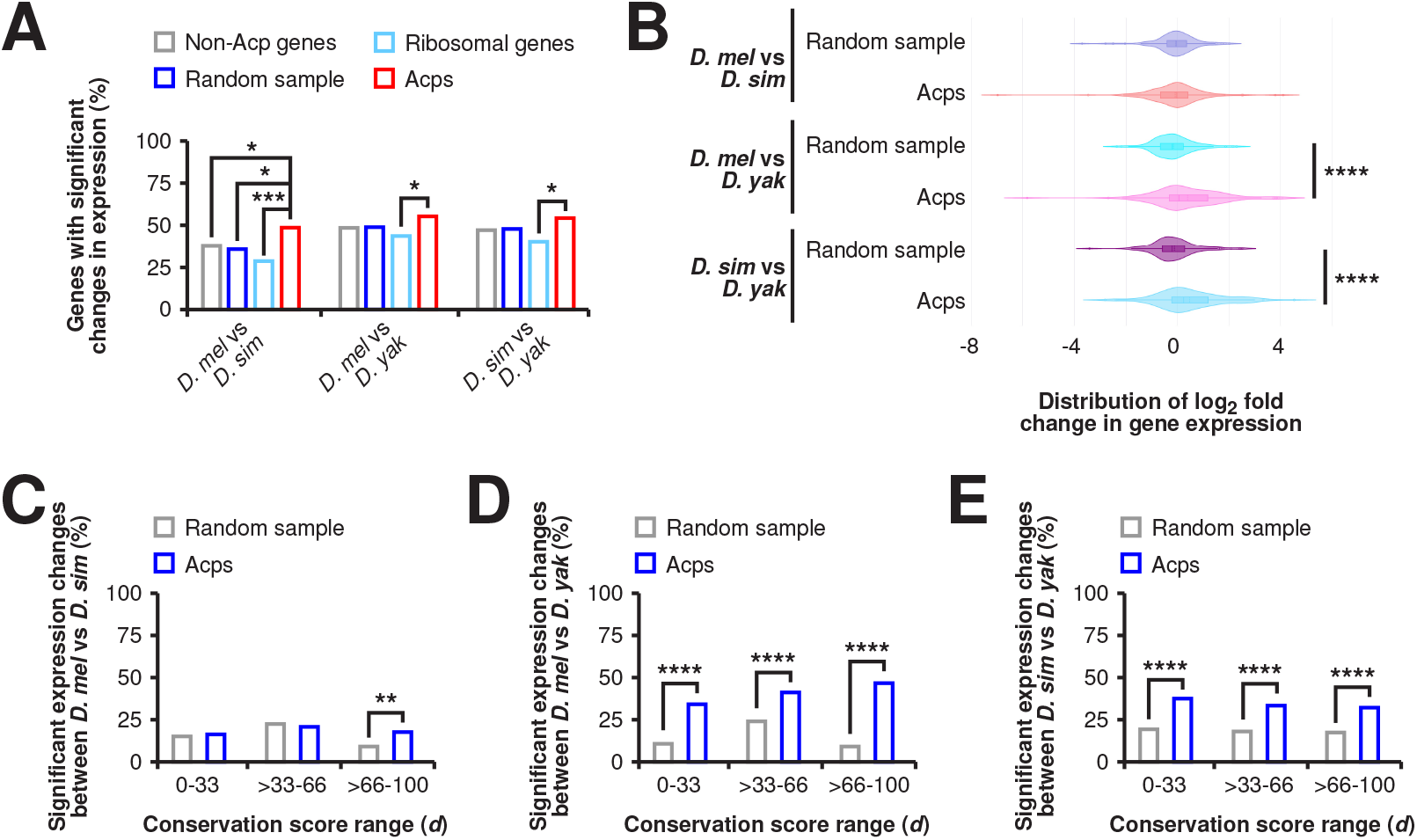
Expression of Acp genes diverges rapidly between closely related *Drosophila* species and correlates with mutation accumulation in promoters. A) The percentage of genes with significant changes in gene expression for Acps (blue) compared to all analysed non-Acp genes (grey) based on differential gene expression analysis between *D. melanogaster* and *D. simulans*, *D. melanogaster* and *D. yakuba*, and *D. simulans* and *D. yakuba*. Statistically significant differences from FDR corrected non-parametric chi-squared tests are indicated by asterisks (* p≤0.05, *** p≤0.001). B) Violin plots of log2 fold changes in gene expression distribution for the random genome sample and Acps based on differential gene expression analysis between *D. melanogaster* and *D. simulans*, *D. melanogaster* and *D. yakuba*, and *D. simulans* and *D. yakuba*. Statistically significant differences from Mann Whitney U tests are indicated by asterisks (**** p≤0.0001). C-E) Comparison of significant changes in log2 fold changes in gene expression in conservation promoter score (*d*) range 0-33%, >33-66%, and >66-100%. Expression changes for Acp (blue) and a random sample of genes were analysed between *D. melanogaster* and *D. simulans*, *D. melanogaster* and *D. yakuba*, and *D. simulans* and *D. yakuba*. Statistically significant differences from non-parametric chi-squared tests are indicated by asterisks (** p≤0.01, **** p≤0.0001).

To analyse whether Acps with rapidly evolving promoters were also differentially expressed between species, we divided Acp and control genes into three conservation score groups based on the calculated *d* score (0-33, >33-66, >66-100 *d*) for each gene and plotted the percentage of genes with significant expression changes for each group (Fig. 2C-E, Supplementary Dataset 1). The random sample control group generally displayed the highest differences in expression for the middle conservation score group (>33-66 *d* score) (Fig. 2C-E). However, Acps were enriched for significant expression changes in the faster evolving group (>66-100 *d* score) across all species comparisons (Fig. 2C-E).

### Acp gene promoters with divergent expression between species display positional hot spots for sequence changes

Next, we further molecularly analysed Acp genes with significant expression changes for promoter evolution in more detail. We selected three representative genes with consistent differences across pairwise species comparisons but with different levels of promoter evolution (*Obp56f*, *CG30486*, and *CG15117*) (Fig. 3A, Supplementary Dataset 1). Further analysis of genes with rapid promoter evolution (*Obp56f* and *CG30486*) revealed substitution hotspots localised to promoters upstream of the TSSs for all three genes (Fig. 3B-D).

**Fig. 3:**
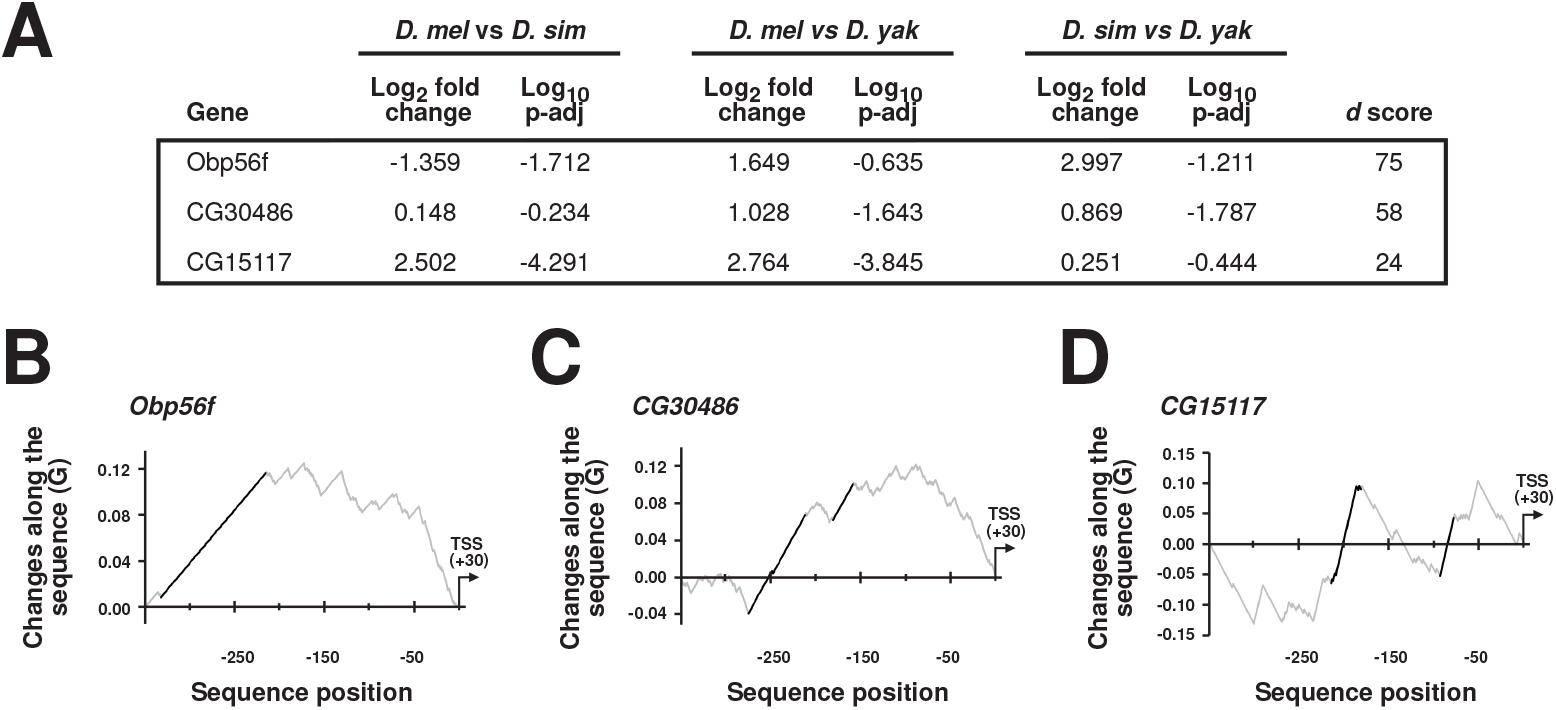
Acp gene promoters with divergent expression between species are hot spots for sequence change accumulation. A) Differential gene expression results for *Obp56f*, *CG30486*, and the control *CG15117* between *D. melanogaster* and *D. simulans*, *D. melanogaster* and *D. yakuba*, and *D simulans* and *D. yakuba*. Significance was considered where log2 fold change in gene expression was ≥0.5 or ≤-0.5, and the adjusted p-value was ≤0.05. B and C) Plots of G scores between nucleotide changes and the differential accumulation of events along fast evolving promoter sequences for *Obp56f* and *CG30486*. Sequences were aligned for *D. melanogaster*, *D. simulans*, *D. sechellia*, *D. yakuba*, and *D. erecta*. Positions in the alignments with significant stretches of substitutions (hot spots) are identified by black lines.

### Divergent expression of Acps with fast evolving promoters is maintained in *Drosophila* hybrids

Interspecies comparisons of differential gene expression cannot rule out the effect of differences in the genomic background and subsequent alteration of the cellular environment that might have arisen during speciation. To test whether changes in the cellular environment of Acp genes can alter expression, we generated hybrids, with a mixed cellular environment, between *D. melanogaster* and *D. simulans* and between *D. melanogaster* and *D. sechellia.* In these hybrids, accessory glands develop normally in contrast to the gonads, thus ruling out possibly allometric effects due to tissue atrophy [62].

To distinguish expression levels of *CG30486*, *Obp56f*, and *CG11598* genes with high promoter evolution in hybrids, we identified single nucleotide polymorphisms (SNPs) that are part of restriction enzyme cut sites. We then designed an RT-PCR amplicon around these SNPs such that PCR fragments of different sizes for each species were obtain after a restriction digest and quantified the levels of PCR products in hybrids (Fig. 4A-D). Analysis of sequence change accumulation hotspots in the *CG11598* promoter indicated a sequence change hot spot (Supplementary Fig. 1). In addition, we also analysed *CG15117* expression in hybrids in the same way because it showed the largest expression changes between these species (Fig. 4E and F). Our analysis revealed that *CG30486* and *CG15117* retain their species-specific expression pattern in hybrids, with both being expressed at higher levels in *D. simulans* and *D. sechellia* compared to *D. melanogaster* (Fig. 4B and F). In contrast, the fastest-evolving promoter genes *Obp56f* and *CG11598* were more expressed in *D. melanogaster* compared to *D. simulans*, but significantly less in *D. melanogaster* compared to *D. sechellia* (Fig. 4C and D). For these two genes, one allele becomes dominant in hybrids. This phenomenon is called transvection and requires chromosome pairing [58].

**Fig. 4:**
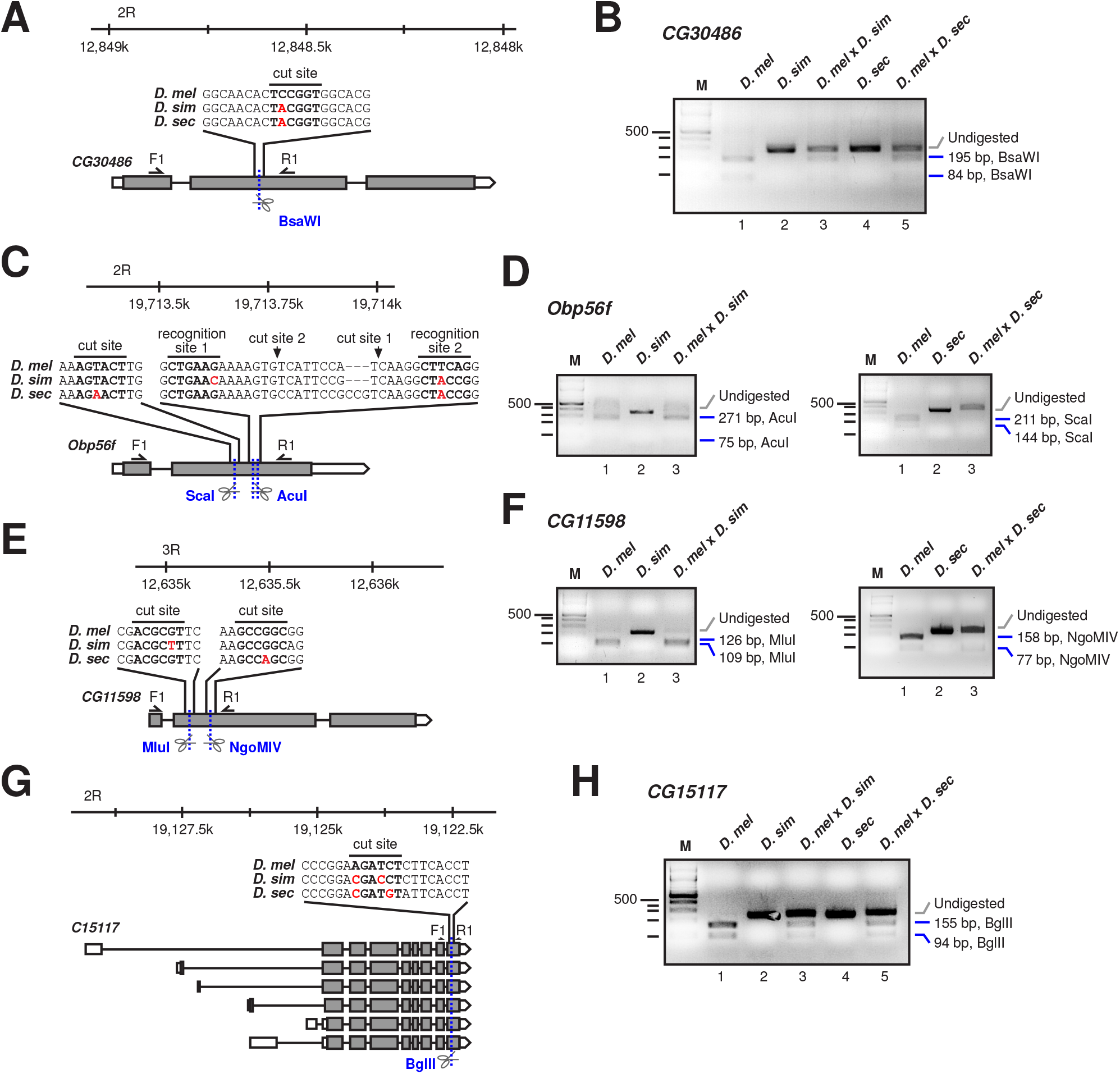
Comparison of transcript expression in *Drosophila* hybrids reveals divergent expression in a subset of Acps with fast evolving promoters. A-H) Schematic representation of the *CG30486* (A), *Obp56f* (C), *CG11598* (E), and *CG15117* (G) gene models and the strategy employed for analysing differential transcript levels in *Drosophila* hybrids. RT PCRs from *D. melanogaster* x *D. simulans*, and *D. melanogaster* x *D. sechellia* hybrids were performed using primers flanking a restriction site with polymorphism(s) either *D. simulans* and/or *D. sechellia*. Primers are indicated as half arrows labelled F1 (forward) and R1 (reverse) and the restriction site alignment is indicated (bold) with polymorphisms in red. The restriction enzymes are also indicated. Cut/recognition sites are indicated in bold and polymorphisms in red. Example agarose gels of *D. melanogaster*-specific restriction enzyme-digested RT PCRs for *CG30486* (B), *Obp56f* (C), *CG11598* (D), and *CG15117* (F). Experiments were performed for *D. melanogaster*, *D. simulans*, *D. melanogaster* x *D. simulans* hybrids, *D. sechellia*, and *D. melanogaster* x *D. sechellia* hybrids. Undigested (*D. simulans* and *D. sechellia*, grey) and digested (*D. melanogaster*, blue) bands are in indicated with sizes and the employed restriction enzymes.

Taken together, our data show that promoter evolution drives changes in expression, because species-specific patterns of expression are either maintained in hybrids or dominated by one allele through transvection. Hence, our data implicate promoter evolution as a main source for fast changes in gene expression.

## Discussion

Acp genes are among the fastest evolving genes. However, the protein coding region of only a minority is under positive selection [24]. Here, we discover that the promoters of Acp genes are hotspots for sequence changes between species. We established a correlation between differences in gene expression among closely related *Drosophila* species and changes occurring in these promoters. Using RT-PCR combined with allele-specific restriction digests in *Drosophila* hybrids, we provide evidence of species-specific divergence in Acp expression driven by rapidly evolving promoters. These findings underscore the critical role of promoter evolution in shaping the diversity of Acp expression across species.

Numerous studies have shown that Sfp gene expression diverges rapidly between related species in both vertebrates and invertebrates [39–44, 63]. Moreover, such changes in expression can be accompanied by functional changes as shown for Sfps in *Lymnaea stagnalis* [63]. Sfps were shown to have divergent gene expression profiles between *D. melanogaster* and *D. simulans* driven by a relaxation of selective pressures [44]. However, Sfps with reproductive-tissue-specific expression were enriched for stabilizing and directional selection, while relaxed selection was the predominant mode of evolution among Sfp genes with other tissue-specific or non-tissue-specific expression [44].

Developmentally redundant Acps can likely tolerate drastic changes in gene expression since perturbation of processes such as sperm competition or the female postmating response do not influence fitness at the organismal level. Expression changes in Acps with postmating-specific roles could drive species divergence and fitness. Here, variability in Sfp concentrations could have drastic changes on reproductive success [44]. For instance, Acps implicated in sperm competition experience high rates of duplication and loss, which could be attributed to increasing or decreasing their expression [6]. Promoter evolution likely has similar implications on expression, which could impact sperm competition. Modifications in the promoters of developmentally vital genes are probably constrained by compensatory mechanisms such as rapidly evolving or multi-choice enhancers [64]. Acps are unlikely restricted by strict stoichiometry from being part of enzymatic complexes, this may explain their general expression divergence [39–44, 63]. As a consequence of changes in expression, however, binding properties likely change. To understand the impact of such changes in expression, we must first learn about the molecular functions of Acps in directing processes such as sperm competition [65, 66].

Of much interest is whether promoter evolution occurs at different rates and what the driving factors behind this are [67]. Notably, it has been shown that promoters of genes required for suppression of transposon mobility evolve rapidly [51–54]. The accumulation of sequence changes in promoter regions has been associated with transposon mobility due to the presence of open chromatin around TSSs [68]. An example of how transposon driven promoter evolution can direct expression changes which affect biological processes is the outcome of *P*-element mutagenesis experiments observed in the *egghead* (*egh*) locus, which encodes an enzyme involved in glycosphingolipid biosynthesis [69]. Following *P*-element mutagenesis, multiple base changes were detected in the first promoter region, leading to the emergence of the SP-insensitive allele, *egh^cm^*. Mutations in the *egh* gene give rise to pleiotropic phenotypes and the *egh^cm^* allele disrupts the neuronal connectivity essential for the female post-mating response and optic lobe development [11, 69, 70].

While testing interspecies expression divergence in hybrids, we observed species-specific gene expression repression for two of the analysed Acps, indicating that one allele becomes dominant. This result is reminiscent of the mechanism of transvection, a unique form of genetic complementation, where regulatory DNA sequences on one homolog can influence the transcriptional activity of a gene on the other homolog as *trans* elements [58, 71]. Transvection requires chromosome pairing, which is the default in *Drosophila* and can result in enhancers activating genes on the opposite homolog (trans-activation), or it can lead to the repression of gene expression across homologous chromosomes (trans-repression) [58]. Interestingly, *cis* elements compete with transvection inducing *trans* elements in a process termed promoter competition [72, 73]. This has been shown in experiments with the promoter-less mutant *Ubx^Mx6^*, which suppresses a haltere-to-wing transformation caused by the presence of a *trans bx^34e^*allele more than a promoter-containing *Ubx^1^* mutant [72, 74]. This suggests that the intronic enhancers in the *Ubx^Mx6^*allele can efficiently *trans*-activate the functional *Ubx* transcription unit on the *bx^34e^* allele [58, 73]. In hybrids, species-specific DNA-based *trans* elements could interact with their species homolog DNA to alter expression by species-specific gene activation or repression. Rapid evolution of promoters in this hybrid setting could reduce promoter competition due to changes in promoter functionality/complementarity caused by sequence evolution.

In summary, our findings show that sequence changes in core promoters present between closely related species result in differences in gene expression. These changes can be maintained in hybrids of closely related species suggesting that promoter evolution dominates over the genomic background that alters cellular environment in hybrids. Moreover, since one allele can become dominant due to transvection in hybrids, our results point out a prominent role for promoter changes in driving evolution.

## Materials and methods

### Defining Acp genes

We based our list of Acp genes on previous published lists [1, 23, 59], and restricted our selection to genes with graded tissue specificity index τ ≥0.9, which was calculated for all genes from FlyAtlas2 RNA expression data [60, 61]. Acp genes with τ ≥0.9, 90% or more of the expression resides in accessory glands. A total of 155 Acps from the original list met these criteria (Supplementary Dataset 1). As a control group, a random sample of 155 genes taken from the genome, excluding the 155 Acps, were defined. Randomisation was performed using the sample function in base R [75].

### Sequence/data retrieval and alignment

Genomic sequences, PhyloP27way data and CAGE data were retrieved from UCSC Genome Browser using the UCSC Table Browser sequence retrieval tool [76, 77]. A standardised sequence region of 2000 bases upstream of the annotated gene TSSs was exported for each gene to ensure inclusion of promoter regions. A region of 1000 nucleotides upstream and 300 nucleotides downstream of gene TSSs were collected from PhyloP27way data. Where genes had multiple TSSs, dominant transcripts were identified using available CAGE values proximal to annotated TSSs. To identify dominant TSSs while accounting for CAGE peak position inaccuracy, a nine-nucleotide sliding window score (*f*) was calculated using R version 4.4.2 for full gene lengths [75]. Annotated TSSs with the highest *f* score per gene were considered dominant and used as representative transcripts for analysis. Sequences were aligned with clustalW using the R package msa and manually refined using the MEGA11 package [78, 79].

### Identification of promotor hot spots and comparison of substitution rates

To analyse promoter evolution hotspots, gene and promoter sequences were adjusted to span -1000/+300 nucleotides from the TSS of each gene. To assess the occurrence of sequence change hotspots within promoter sequences, we computed the hotspot accumulation score (*d*) using alignments derived from the trimming process, or alternatively, PhyloP data. These alignments were converted into ’events’ for each species when compared to *D. melanogaster*. Here, each nucleotide in the sequence was assigned a value of 0 if it represented a conserved sequence and 1 if it indicated a sequence change (either a base change or an indel event). Events were tallied for all changes, as well as separately for base changes and indels. The cumulative events were calculated for both concatenated gene groups and individual genes across all nucleotides. Subsequently, a sliding event (*Se*) score was computed from these tallies using a sliding window of five bases along the sequence. To determine the percentage of events surpassing the average control promoter *Se* score (*d*), a 350-nucleotide region upstream of the estimated TATA box region was examined. This involved dividing the total number of *Se* scores exceeding the average sliding event score of the control group (*Se^C^*) by the total number of events within that region (*N*). To compute the promoter region scores (*d^P^*) from PhyloP data, we substituted the total number of PhyloP (*p*) scores in the 350-nucleotide regions that were below the average of the control group promoter region (*p^C^*) in place of the total number of *Se* events where *Se* was greater than *Se^C^* as previously described. Significance was assessed using non-parametric chi-squared tests compared to the control group *d* score. Significance values where p≤0.05 after Bonferroni correction were considered statistically significant.

### Accumulation of substitutions along extended gene regions

To test for nonrandom accumulation of indels along the Acp extended gene regions between the five analysed *Drosophila* species (*D. melanogaster*, *D. simulans*, *D. sechellia*, *D. yakuba*, and *D. erecta*) we determined significant deviations from a uniform distribution of substitutions using an empirical cumulative distribution function [80]. The position of the indel event was defined as the 5’ site of the start of the indel in the alignment as per [81]. The function (G) detects monotonic increases in substitutions (n) measured as the difference between the relative occurrence of a nucleotide change and its relative position in the alignment [80]. Whether differences between the values of the G function (ΔG) between substitutional events deviates from a random accumulation of changes is tested using Monte Carlo simulations to produce 100,000 samples of n events by sampling sites without replacement along the alignment [80].

### Comparative gene expression analysis between *D. melanogaster*, *D. simulans* and *D. yakuba*

We utilised publicly available RNA-seq data from *D. melanogaster*, *D. simulans*, and *D. yakuba* male whole flies for our analysis (GSE28078) [82]. Adapter sequences were removed from the raw reads using the Trimmomatic tool (v0.39) with default parameters. Trimmed reads were aligned to the respective genome assemblies (dm6 for *D. melanogaster*, Prin_Dsim_3.1 for *D. simulans*, and droYak2 for *D. yakuba*) using the RNA-seq aligner RNA STAR (v2.7.10b). Gene counts were quantified using the featureCounts tool from the Subread package (v2.0.3). Orthologous genes were identified using the Flybase ftp accession orthologue lookup table [83]. Following this, gene count normalization and the differential gene expression analysis between the two species were conducted employing DESeq2 (v2.11.40.8) with an additional step for species-specific gene length normalisation. Analysis was performed using usegalaxy.eu [84].

### *Drosophila* stocks, genetics, and hybrid culture

Wild type *D. melanogaster* CantonS, *D. simulans* and *D. sechellia* Jallons (S-32) were used and kept on standard cornmeal-agar food (1% industrial-grade agar, 2.1% dried yeast, 8.6% dextrose, 9.7% cornmeal and 0.25% Nipagin, all in (w/v)) in a 12 h light:12 h dark cycle. To obtain hybrid males, *D. melanogaster* males were crossed to either *D. simulans* or *D. sechellia* virgin females.

### RNA extraction and RT-PCR

Total RNA extraction was carried out using Tri-reagent from SIGMA according to the manufacturer’s instructions. Reverse transcription was performed using Superscript II (Invitrogen) using an oligo-dT17V primer as previously described (Haussmann et al Genetics, 2011). PCR for *CG30486*, *Obp56f* and *CG11598* was performed for 40 cycles with 1 μl of cDNA. Primers used are listed in the Supplementary Dataset 1. Restriction enzymes (BsaWI, AcuI, MluI, ScaI, or NgoMIV) were from New England Biolabs and DNA restriction digests were performed according to manufacturer’s instructions. Experiments included at least two biological replicates.

## Supplementary information

Supplementary information includes Supplementary Dataset S1.

## Acknowledgements

We thank C. Rezaval and S. Collier for *Drosophila* species, and the Soller lab for helpful discussion. *Drosophila* images were acquired from D. J. Obbard (https://obbard.bio.ed.ac.uk).

## Authors’ contributions

D.W.J.M. and M.S. conceived the project. D.W.J.M. directed the project. D.W.J.M. and F.H.S.S. carried out evolutionary analysis. D.W.J.M. performed genetic experiments and molecular biology. R.A., D.W.J.M. and A.F. carried out RNA-seq analysis. D.W.J.M. wrote the manuscript and M.S. and A.C reviewed it. All authors read and approved the final manuscript.

## Availability of data and materials

All data generated or analysed during this study are included in the supplementary information files.

## Declarations

### Ethics approval and consent to participate

Not applicable.

## Consent for publication

Not applicable.

