## Supplementary Figure 1 for "Rapid promoter evolution of male accessory gland genes is accompanied by divergent expression in closely related *Drosophila* species"

**Supplementary Fig. 1: Hot spots for sequence change accumulation in the Acp gene *CG11598* promoter**

Plot of G scores between nucleotide changes and the differential accumulation of events along the fast-evolving *CG11598* promoter sequence. Sequences were aligned for *D. melanogaster*, *D. simulans*, *D. sechellia*, *D. yakuba*, and *D. erecta*. Positions in the alignment with significant stretches of substitutions (hot spots) are identified by black lines.

### Supplementary Figure 1

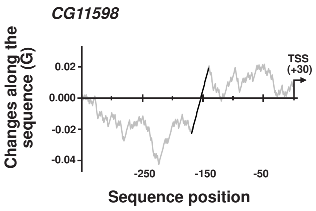
